# Pan-resistant *Candida auris*: New York Sub-cluster Susceptible to Antifungal Combinations

**DOI:** 10.1101/2020.06.08.136408

**Authors:** Brittany O’Brien, Jiali Liang, Sudha Chaturvedi, Jonathan L. Jacobs, Vishnu Chaturvedi

## Abstract

Four pan-resistant *Candia auris* strains from New York outbreak were 100% inhibited *in vitro* by combinations of two antifungal drugs using fixed concentrations achievable *in vivo*. Pan-resistant *C. auris* strains have mutations in eleven gene targets associated with major antifungal drugs, and constituted a distinct sub-cluster among NY strains.

---

We recently reported the emergence of pan-resistance in *Candida auris* from New York ^1^. Since 2016, New York hospitals and healthcare facilities have faced the highest number of clinical and surveillance cases of *C. auris* in the United States^2^. Effective strategies for the prevention, control, and treatment of *C. auris* are still being developed, all of which could be complicated by the observed pan-resistance. A conceptual framework supports deploying drug combinations to combat the threat of antimicrobial resistance^3^. Therefore, we studied if strains of pan-resistant *C. auris* are susceptible to combinations of current antifungal drugs and what genetic features distinguish NY pan-resistant *C. auris*. Details of methods are provided in the Supplementary Appendix. Four pan-resistant *C. auris* strains were 100% inhibited *in vitro* by combinations of two antifungal drugs using fixed concentrations achievable *in vivo*. Expectedly, flucytosine combinations with amphotericin B, azoles, or echinocandins were most effective (Figure 1A). In time-kill analysis, two-drug combinations caused greater than two log reduction in growth relative to the respective single drugs suggestive of a fungicidal action (Figure 1B). These results are consistent with our recent publication on the efficacy of antifungal combinations for NY *C. auris* strains with various multi-drug resistance patterns. Based on a comparative genomic analysis, we found four pan-resistant *C. auris* strains have mutations in eleven gene targets associated with major antifungal drugs (Figure 1C). These findings are similar to other reports for drug-resistant *C. auris* strains. Two different, non-synonymous mutations in the predicted sequence of FKS1p were observed. *C. auris* 19-4 and 19-61 exhibited FKS1 S635P, whereas *C. auris* 19-42 and 19-43 showed FKS1 S635Y (Figure 1D).

**Figure 1.**
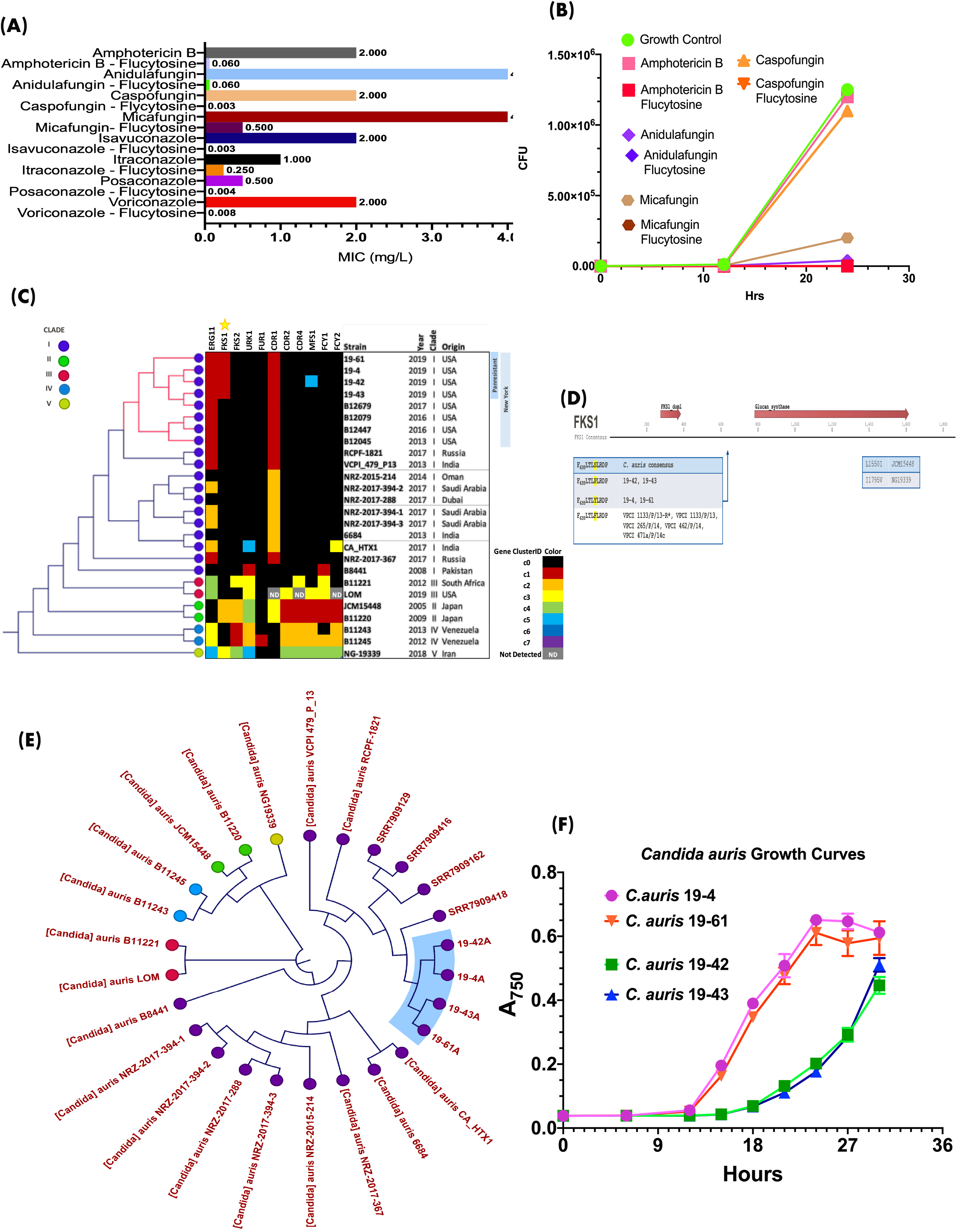
Characterization of pan-resistant *Candida auris*. (A) Pan-resistant *C. auris* strains are susceptible to two-drug combinations containing either amphotericin B, an azole or an echinocandin in combination with flucytosine; representative data shown for *C. auris* 19-4, (B) Time-kill curve with single drug alone or two-drug combination with flucytosine. Two-drug combinations caused more than two log reduction in growth; representative data shown for *C. auris* 19-43, (C) Four pan-resistant *C. auris* strains (19-4, 19-42, 19-43, 19-61) have gene variants associated with resistance to a variety of antifungal drugs, (D) Notably, mutations in the hotspot of glucan synthase gene FKS1, the target of echinocandins are found in *C. auris* 19-4, 19-42, 19-43, and 19-61, (E) Four pan-resistant *C. auris* strains constituted a distinct sub-cluster among NY strains; the unrooted Neighbor-Joining tree was derived from whole genome assemblies, (F) A profound lag in growth of two pan-resistant strains, *C. auris* 19-42 and *C. auris* 19-43 suggesting a fitness cost; RPMI broth at 35°C.

These mutations are in a known hotspot of FKS1, a glucan synthase gene, and the target of echinocandin antifungal drugs. All four pan-resistant strains constituted a distinct sub-cluster among NY strains (Figure 1E) ^4,5^. Finally, pan-resistance appears to exact fitness cost in at least two *C. auris* strains (19-42 and 19-43), which showed a prolonged lag growth phase (Figure 1F); these strains have a high resistance to caspofungin (>16 mg/L). Further results are presented in the Supplementary Appendix. Our findings suggest that pan-resistant *C. auris* strains remain susceptible to antifungal combinations, which might help expand the available therapeutic options. The genetic analysis suggests ongoing mutations in response to antifungal drug pressure are the likely drivers of emerging pan-resistance in NY *C. auris* strains.

We thank valuable assistance of YanChun Zhu, Lynn Leach, and MD. Rokebul Anwor (Mycology Laboratory), Matt Shudt (Applied Genomic Technologies Core), and Jonathan Adams (Media and Glassware Core) Wadsworth Center, New York State Department of Health, Albany, NY, USA. We also acknowledge continuing efforts of *C. auris* Investigation Workgroup, Division of Epidemiology, New York State Department of Health, Albany, NY, USA. Supported in part by a Cooperative Agreement number NU50CK000516, funded by the Centers for Disease Control and Prevention. Its contents are solely the responsibility of the authors and do not necessarily represent the official views of the Centers for Disease Control and Prevention or the Department of Health and Human Services.

