## Supplementary data for "Pan-resistant *Candida auris*: New York Sub-cluster Susceptible to Antifungal Combinations"

**Supplementary Appendix**

Supplementary Methods	2-5

Acknowledgements	6

Supplementary Tables	7-11

Supplementary Figures	12-21

References	22

**Supplementary Methods**

*Candida auris* characterization and antifungal testing: *C. auris* surveillance and clinical samples were tested using selective culture media, and fungal isolates were identified by mass spectrometry (MALDI-TOF-MS), and Sanger sequencing of ribosomal ITS2 genes ^1,2^. Conventional antifungal susceptibility testing with single drugs was according to the CLSI M-60A method and susceptibility breakpoints interpreted per CDC recommendations ^3^. The metadata and antifungal susceptibility test results are summarized in supplementary table 1. Fixed concentration of two-drug combinations was tested in custom-made plates from Trek Diagnostics System and 100% inhibition end-points recorded per recent publication by our group ^4^. The results of two-drug combination tests are summarized in supplementary 2. There are no approved methods for time-kill (fungicidal) testing of antifungal drugs; we adapted essential parameters of an approved method for bactericidal activities of antimicrobial agents ^5^. We used 96-well microtiter plates instead of glass tubes. All drugs stock solutions were prepared in DMSO (Sigma-Aldrich, St. Louis, MO, USA). For inoculum, the initial cell suspension in RPMI-1640 was adjusted to absorbance (A_530_) of 0.08 to 0.1, 20 μl suspension added to 11 ml of RPMI 1640 broth, and 100 μl of the final suspension added to each well of the 96-well microtiter plate^4^. Duplicate wells containing growth control, single drugs, and drug combinations with flucytosine at fixed concentrations, were inoculated with each test isolate, and sampled at 0, 6, 12, and 24 hours by withdrawing 100 µl cell suspension. After 10-fold serial dilutions of the cell suspension in sterile physiological saline, the cell suspensions were plated on Sabouraud Dextrose Agar plates and incubated at 35℃ for 48 hours. Colony forming units (CFU) determined from the average of readings obtained from two adjacent wells. Two log or higher reduction (≥99.3%) in CFU of treated well vis-à-vis growth control was considered as minimum fungicidal concentration (MFC) as shown in the supplementary table 3 and supplementary figure 1.

Fitness of four *C. auris* isolates was tested by monitoring their growth in RPMI-1640 medium, in stationary cultures in 96-well microtiter plates at 35 ℃, and A_750_ recorded at periodic intervals with a BioTek Synergy 2 plate reader. Data were analyzed and plotted using GraphPad Prism 8 for Mac. We examined cellular aggregation among pan-resistant *C. auris* isolates per an earlier publication ^6^; no notable aggregation was observed.

*Candida auris* genome sequencing: Fungal DNA for genome sequencing was prepared using Qiagen QIAamp DNA mini-kit, quantities measured with Qubit and Nanodrop, and submitted to the Wadsworth Center Applied Genomic Technologies Core for sequencing using Nextera Flex library, 2x250 bp sequencing run on an Illumina NexSeq instrument with an expected output of 100 X or higher genome coverage ^7^. *Candida auris* genome sequence data generated have been submitted to the SRA as BioProject ID PRJNA640677.

Bioinformatics Analysis of *Candida auris* isolates: Following sequencing of each isolate, FASTQ files were imported into CLC Genomics Workbench v.20 with the Microbial Genomics Module software (QIAGEN A/S, Denmark). Except where otherwise noted, all downstream bioinformatics analysis was performed with CLC. For comparative genomics purposes, additional whole genome sequencing data was downloaded from the NCBI’s Sequence Read Archive (REFS). Draft assemblies of each panresistant isolate were constructed using CLC Genomics Workbench v20. In addition, assemblies were constructed for isolates where raw sequencing data was available in NCBI’s Sequence Read Archive, but for which no assembly was available (B12679, B12079, B12447, B12045, and NG-19339). Assembly details are available in supplementary table 4, and average nucleotide identity of the assembled genomes is shown in supplementary table 5.

Methods of Phylogenetic Reconstruction: The whole genome alignment of *C. auris* from available sequences in public database is shown in figure 2. A phylogenetic *k*-mer tree was created using whole-genome draft assemblies of each pan-resistant isolate, as well as multiple *C. auris* reference assemblies representing all known clades (supplementary figure 3). The tree was created using an implementation of feature frequency profile (FFP) via Jensen-Shannon divergences as implemented in QIAGEN CLC Microbial Genomics Module. A *k*-mer length of 16, and only those with the prefix ATGAC on either strand was selected. *C. auris* SNP tree created in CLC Genomics Workbench with Microbial Genomics Module, is shown as supplementary figure 4. FASTQ files from the four pan-resistant strains (19-4, 19-42, 19-43, and 19-61), as well as raw data obtained from NCBI Sequence Read Archive (SRR3883466, SRR7909129, SRR7909416, SRR7909418, SRR10290621, SRR10277313, SRR10292311, SRR10290272, SRR10292114, SRR10292063, ERR899743, SRR9201318, SRR1664626, SRR10851769, SRR3883453, SRR3883452, DRR129819, SRR3883464, and SRR9007776) was mapped to the B8441 reference assembly. Read mappings and variant tracks were used to construct a SNP-tree, where SNPs were filtered such that minimum coverage required in each sample at each position was >= 30, and individual SNPs required 25% depth of coverage relative to each whole genome mapping. SNPs closer than 10 nt distant were pruned to reduce complexity, and only those SNPs with a p-value <= 0.01 were considered. Maximum likelihood estimation with a general time reversible nucleotide substitution model, and 1,000 bootstrap replicates was use for tree construction. Branches with confidence intervals above 80% are highlighted in bold.

Resistance Gene Analysis: NCBI protein databases were searched for genes associated with resistance to each antifungal drug tested using free text queries, such as “(fungi[filter] AND "Candida"[All Fields]) OR (txid498019[Organism:noexp]) ) AND “Gene Name”[all fields]”. The resulting protein sequence lists were downloaded and manually curated to remove partial sequencing results or unrelated proteins. The final list of protein homologs for each gene were subjected to multiple sequence alignment to construct a consensus sequence. This consensus was then used as a query with tblastn to identify the closest homolog in each of the panresistant isolates. Protein sequences from each BLAST search were extracted and the multiple sequence alignment was recalculated for each family of resistance proteins. Non-synonymous mutations were identified manually, and phylogenetic trees were created for each protein family using a neighbor-joining algorithm with a Jukes-Cantor distance metric and 1,000 bootstrap replicates. Clusters of related proteins from each homolog family were assigned manually and reported in Figure 1B as “cluster IDs”. Individual trees for each protein family are also shown (Supplementary figures 5-14). All analysis was carried out in CLC Genomics Workbench v20. We also compared identified antifungal gene targets with the earlier published reports on various drug resistance genes in *C. auris* ^3,8-11^

Acknowledgments

We thank valuable assistance of YanChun Zhu, Lynn Leach, and MD. Rokebul Anwor (Mycology Laboratory), Matt Shudt (Applied Genomic Technologies Core), and Jonathan Adams (Media and Glassware Core) Wadsworth Center, New York State Department of Health, Albany, NY, USA. We also acknowledge continuing efforts of C. auris Investigation Workgroup, Division of Epidemiology, New York State Department of Health, Albany, NY, USA

Supplementary Table 1. Characteristics of pan-resistant *C. auris* isolates and antifungal susceptibility test results

| Isolate | Sample type | Source | Clade | Fluconazole | Voriconazole | Itraconazole | Isavuconazole | Posaconazole | Anidulafungin | Caspofungin | Micafungin | Amphotericin B |
| --- | --- | --- | --- | --- | --- | --- | --- | --- | --- | --- | --- | --- |
| 19-4 | Isolate | Urine | Clade I | >256**^#^** | 2 | 0.5 | 0.5 | 0.25 | 4 | 2 | 4 | 2* |
| 19-42 | Isolate | Blood | Clade I | >256 | 2 | 1 | 1 | 0.25 | 4 | 16 | 4 | 2 |
| 19-43 | Isolate | Blood | Clade I | >256 | 2 | 1 | 1 | 0.25 | 4 | 16 | 4 | 2 |
| 19-61 | Isolate | Urine | Clade I | >256 | 2 | 0.5 | 0.5 | 0.25 | 4 | 2 | 4 | 2 |

**^#^** antifungal susceptibility testing was performed using CLSI M-60 methods; results are mg/L.

**^*^** amphotericin B was tested using E-test strips; more details in our recent publications^2^

Supplementary Table 2. *In vitro* susceptibility of pan-resistant *Candida auris* isolates to fixed concentrations of two-drug antifungal combinations*

|  | *C. auris* 19-4 | *C. auris* 19-61 | *C. auris* 19-42 | *C. auris* 19-43 |
| --- | --- | --- | --- | --- |
| Amphotericin B (AMB) | 1**^#^** | 1 | 1 | 0.5 |
| 5-flucytosien (FLC) | 0.5 | 0.25 | 0.125 | 0.125 |
| AMB/FLC | 0.06/0.25 | 0.0625/0.25 | 0.03125/0.125 | 0.0078/0.5 |
| Anidulafungin (AFG) | >4 | >4 | 4 | >4 |
| AFG/FLC | 0.06/0.5 | 0.0624/0.5 | 0.0624/0.4992 | 0.0624/0.5 |
| Caspofungin (CAS) | >4 | >4 | >4 | >4 |
| CAS/FLC | 0.0039/0.4992 | 0.0039/0.5 | 0.0039/0.4992 | 0.0039/0.5 |
| Micafungin (MFG) | >4 | >4 | >4 | >4 |
| MFG/FLC | 0.5/0.25 | 0.5/0.25 | 0.5/0.25 | 0.25/0.125 |
| Isavuconazole (ISA) | >2 | 2 | 2 | >2 |
| ISA/FLC | 0.0039.0.5 | 0.0039/0.5 | 0.0039/0.4992 | 0.0039/0.5 |
| Itraconazole (ITC) | 1 | 1 | 0.5 | 0.5 |
| ITC/FLC | 0.25/0.5 | 0.0625/0.125 | 0.125/0.25 | 0.0625/0.125 |
| Posaconazole (POS) | 0.5 | 0.5 | 0.5 | 0.5 |
| POS/FLC | 0.0039/0.4992 | 0.0039/0.5 | 0.0039/0.4992 | 0.0039/0.5 |
| Voriconazole (VOR) | >2 | >2 | >2 | >2 |
| VOR/FLC | 0.0078/0.4992 | 0.0078/0.5 | 0.0078/0.4992 | 0.0078/0.5 |

*Drug concentration (mg/L) required for one-hundred percent inhibition as detailed in a recent publication^4^

\# MIC readings for AMB are lower in broth assay than E-test results in table 1

Supplementary Table 3. Time-kill testing of panresistant *C. auris* isolates against two-drug combination of antifungals (24-hr readings shown)

| Isolate | Drug | Drug Conc.**^#^** | Mean CFU Growth Control**^*^** | Mean CFU with Drug (s) | Mean %inhibition | Log difference |
| --- | --- | --- | --- | --- | --- | --- |
| 19-4 | AMB | 2 | 4,350,000 | 4,200,000 | 3.45% | 0.015 |
|  | AMB+FLC | 0.5 | 4,350,000 | 9,600 | 99.78% | 2.656 |
|  | CAS | 2 | 4,350,000 | 3,600,000 | 17.24% | 0.082 |
|  | CAS+FLC | 0.5 | 4,350,000 | 1,1800 | 99.73% | 2.567 |
|  | AND | 4 | 4,350,000 | 135,000 | 96.90% | 1.508 |
|  | AND+FLC | 0.5 | 4,350,000 | 9,550 | 99.78% | 2.658 |
|  | MFC | 4 | 4,350,000 | 250,000 | 94.25% | 1.241 |
|  | MFC+FLC | 0.5 | 4,350,000 | 8,300 | 99.81% | 2.719 |
| 19-42 | AMB | 2 | 3,450,000 | 2,900,000 | 15.94% | 0.075 |
|  | AMB+FLC | 0.5 | 3,450,000 | 1,950 | 99.94% | 3.248 |
|  | CAS | 2 | 3,450,000 | 2,800,000 | 18.84% | 0.091 |
|  | CAS+FLC | 0.5 | 3,450,000 | 800 | 99.98% | 3.635 |
|  | AND | 4 | 3,450,000 | 35,000 | 98.99% | 1.994 |
|  | AND+FLC | 0.5 | 3,450,000 | 1,400 | 99.96% | 3.392 |
|  | MFC | 4 | 3,450,000 | 1,050,000 | 69.57% | 0.517 |
|  | MFC+FLC | 0.5 | 3,450,000 | 550 | 99.98% | 3.797 |
| 19-43 | AMB | 2 | 2,550,000 | 3,300,000 | 0.00% | 0.000 |
|  | AMB+FLC | 0.5 | 2,550,000 | 1,550 | 99.94% | 3.216 |
|  | CAS | 2 | 2,550,000 | 2,650,000 | 0.00% | 0.000 |
|  | CAS+FLC | 0.5 | 2,550,000 | 1,050 | 99.96% | 3.385 |
|  | AND | 4 | 2,550,000 | 20,000 | 99.22% | 2.106 |
|  | AND+FLC | 0.5 | 2,550,000 | 1,100 | 99.96% | 3.365 |
|  | MFC | 4 | 2,550,000 | 1,050,000 | 58.82% | 0.385 |
|  | MFC+FLC | 0.5 | 2,550,000 | 1,700 | 99.93% | 3.176 |
| 19-61 | AMB | 2 | 3,950,000 | 6,000,000 | 0.00% | 0.000 |
|  | AMB+FLC | 0.5 | 3,950,000 | 19,800 | 99.50% | 2.300 |
|  | CAS | 2 | 3,950,000 | 2,350,000 | 40.51% | 0.226 |
|  | CAS+FLC | 0.5 | 3,950,000 | 22,800 | 99.42% | 2.239 |
|  | AND | 4 | 3,950,000 | 505,000 | 87.22% | 0.893 |
|  | AND+FLC | 0.5 | 3,950,000 | 23,800 | 99.40% | 2.220 |
|  | MFC | 4 | 3,950,000 | 1,250,000 | 68.35% | 0.500 |
|  | MFC+FLC | 0.5 | 3,950,000 | 17,800 | 99.55% | 2.346 |

**^#^**mg/L

**^*^**CFU, colony forming unit; the inoculum comprised of approximately 1,000 CFU

Supplementary Table 4. Metadata for pan-resistant *C. auris* genomes

| **NGS Library** | **19-42** | **19-43** | **19-4** | **19-61** |
| --- | --- | --- | --- | --- |
| **Reads** | **10,661,204** | **9,053,680** | **10,453,020** | **10,054,084** |
| **Bases** | **2,565,429,260** | **2,163,252,185** | **2,499,913,980** | **2,399,436,924** |
| **Trimmed Reads** | **10,652,413** | **9,044,141** | **10,443,215** | **10,044,270** |
| **DeNovo Assembly** |  |  |  |  |
| **Mapped Reads** | **10,600,121** | **9,002,227** | **10,380,806** | **9,998,748** |
| **Contigs** | **723** | **584** | **876** | **637** |
| **ReMappedReads** | **10,604,913** | **9,004,572** | **10,384,230** | **10,001,281** |
| **Refined Contigs** | **648** | **503** | **806** | **571** |
| **Reduction%** | **10.4%** | **13.9%** | **8.0%** | **10.4%** |
| **Genome Size Est. (MB)** | **12.59** | **12.53** | **12.51** | **12.51** |

Supplementary Table 5: Average Nucleotide Identity (ANI) and alignment percentages of four pan-resistant *C. auris* isolates relative to other genomes available in the database.

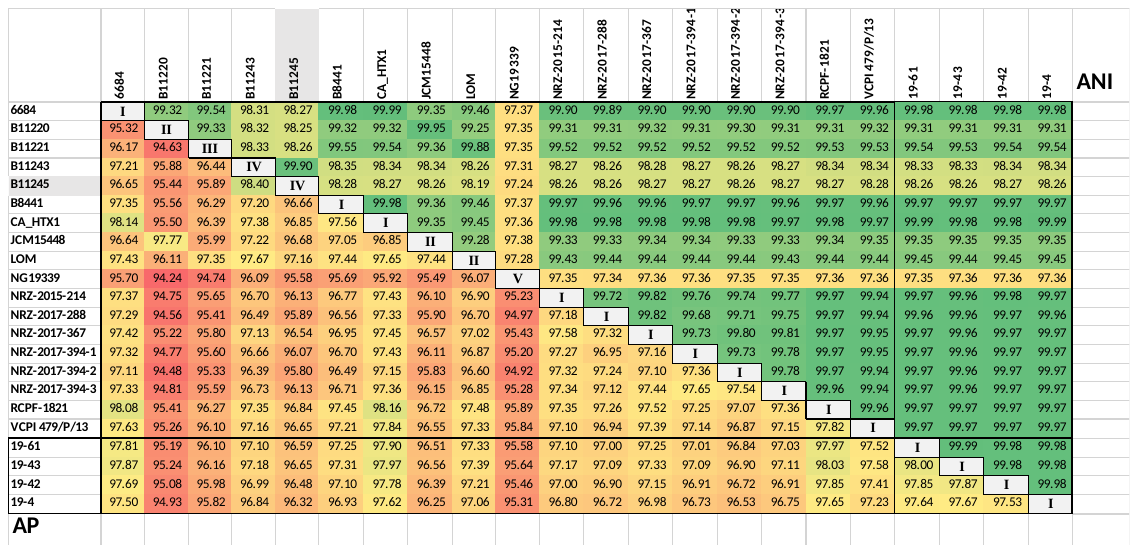

Average nucleotide identity (ANI, upper) and alignment percentage (AP, lower) of comparing whole genome assemblies representing all major reference isolates and clades against one another.

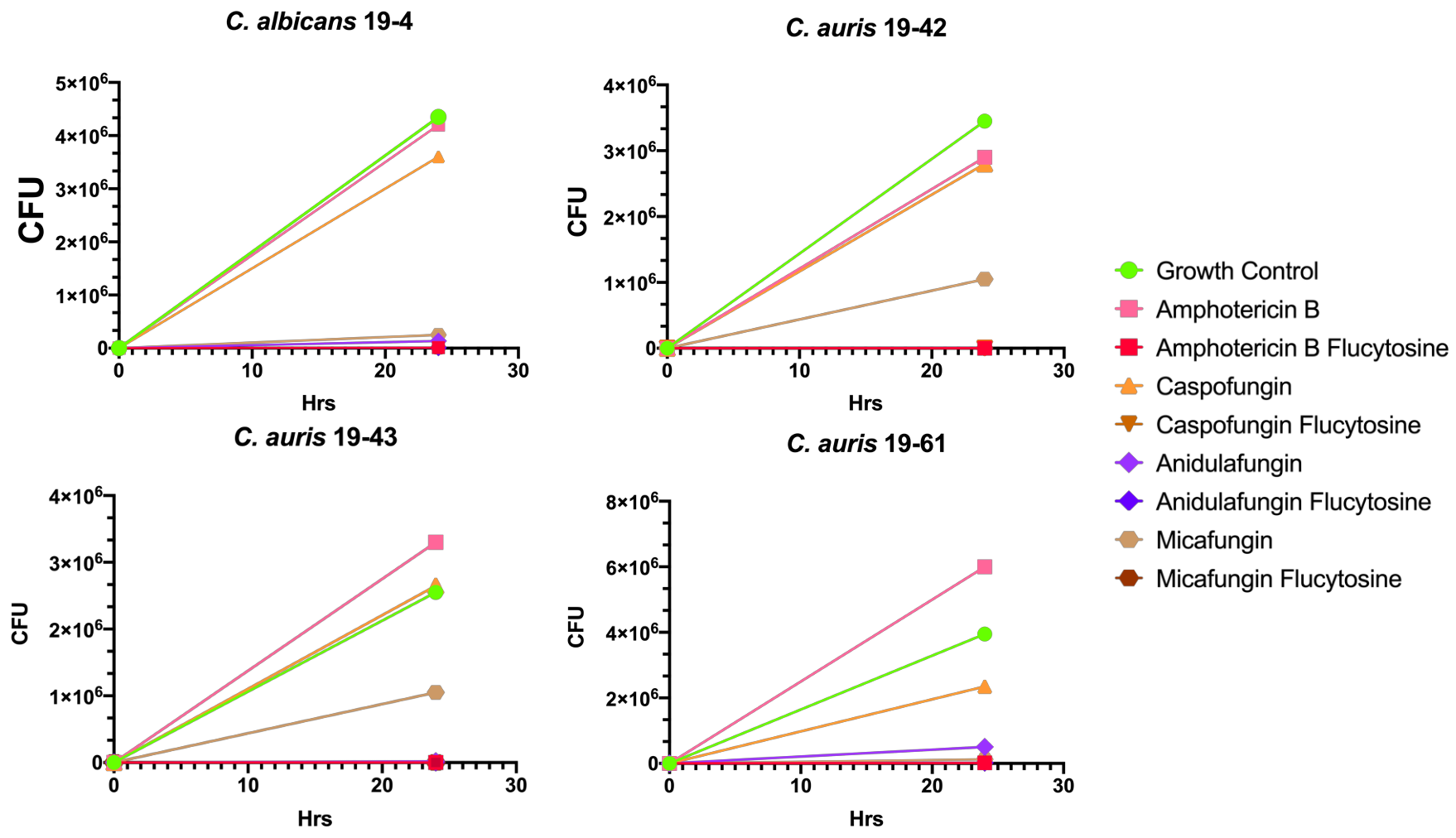
**Supplementary Figure 1**

**Supplementary Figure 2**

Whole Genome Alignment of *Candida auris* from available sequences in public database.

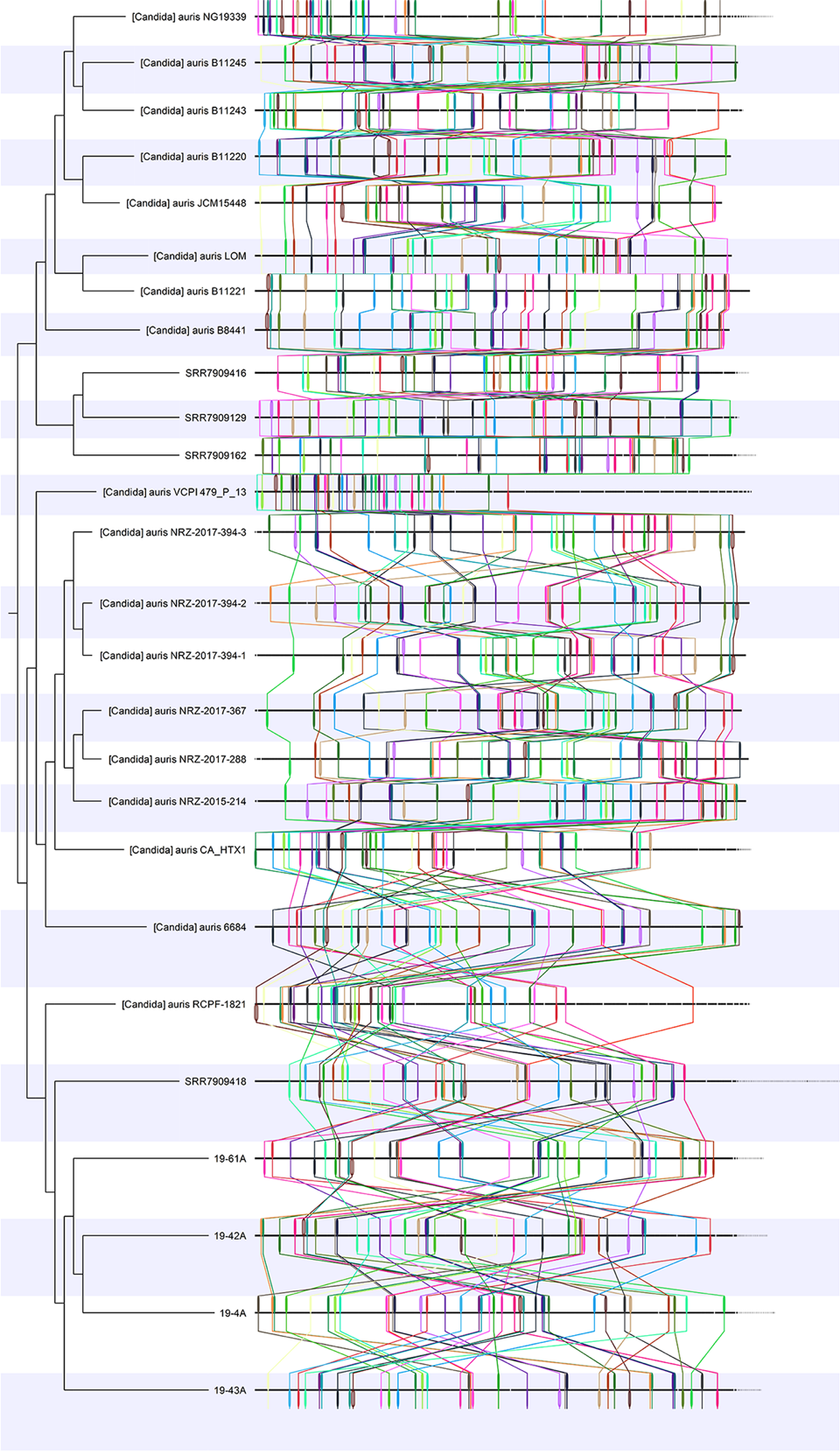

**Supplementary Figure 3**

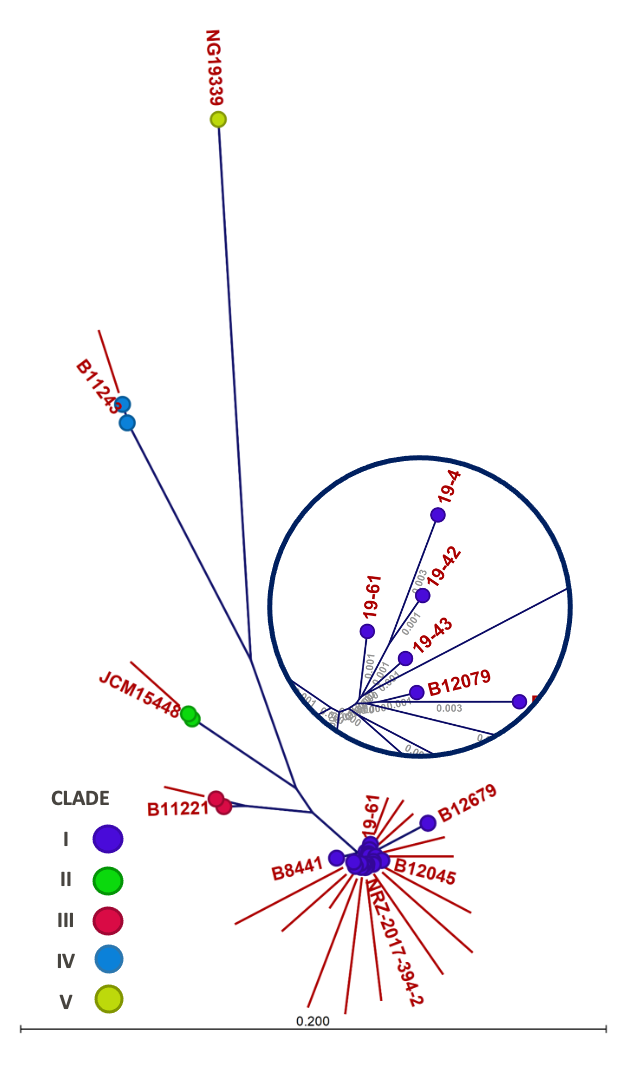
A phylogenetic k-mer tree was created using whole-genome draft assemblies of each panresistant isolate, as well as multiple *Candida auris* reference assemblies representing all known Clades. The list of reference strains and their accession numbers from NCBI is shown in Table 1. The tree was created using an implementation of feature frequency profile (FFP) via Jensen-Shannon divergences as implemented in QIAGEN CLC Microbial Genomics Module. A k-mer length of 16, and only those with the prefix ATGAC on either strand was selected**.**

**Supplementary Figure 4**

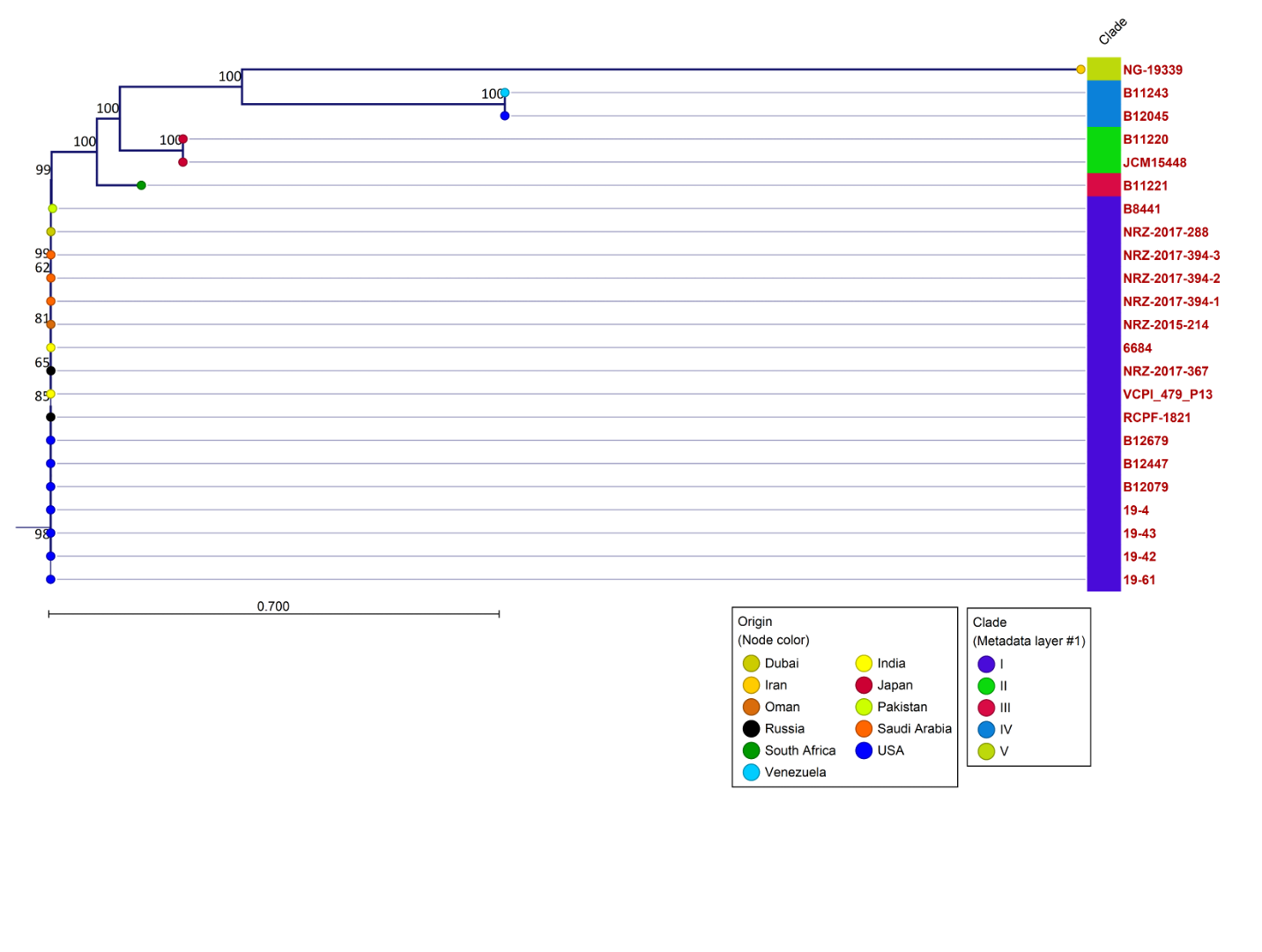
*C. auris* SNP tree created in CLC Genomics Workbench with Microbial Genomics Module, maximum likelihood estimation with a general time reversible nucleotide substitution model, and 1,000 bootstrap replicates (details in text

**upplementary Figures 5-6**

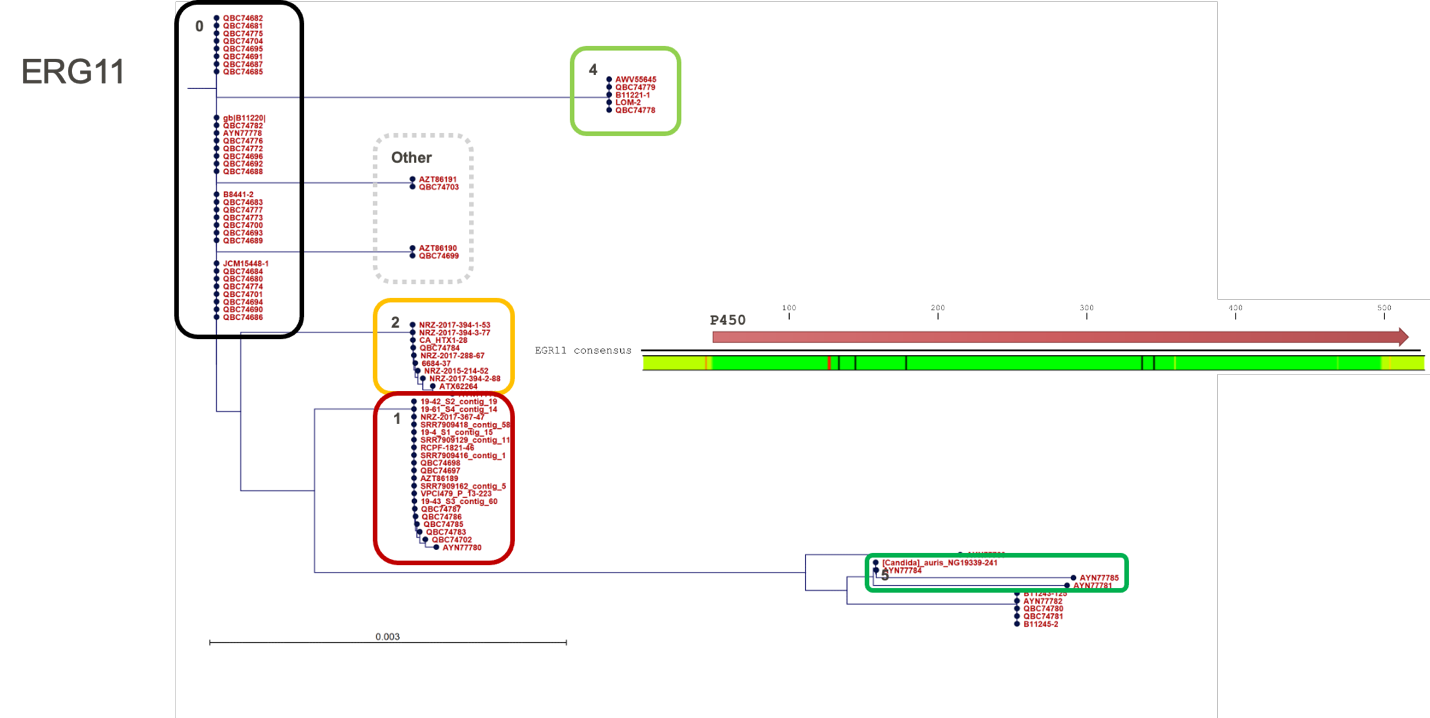

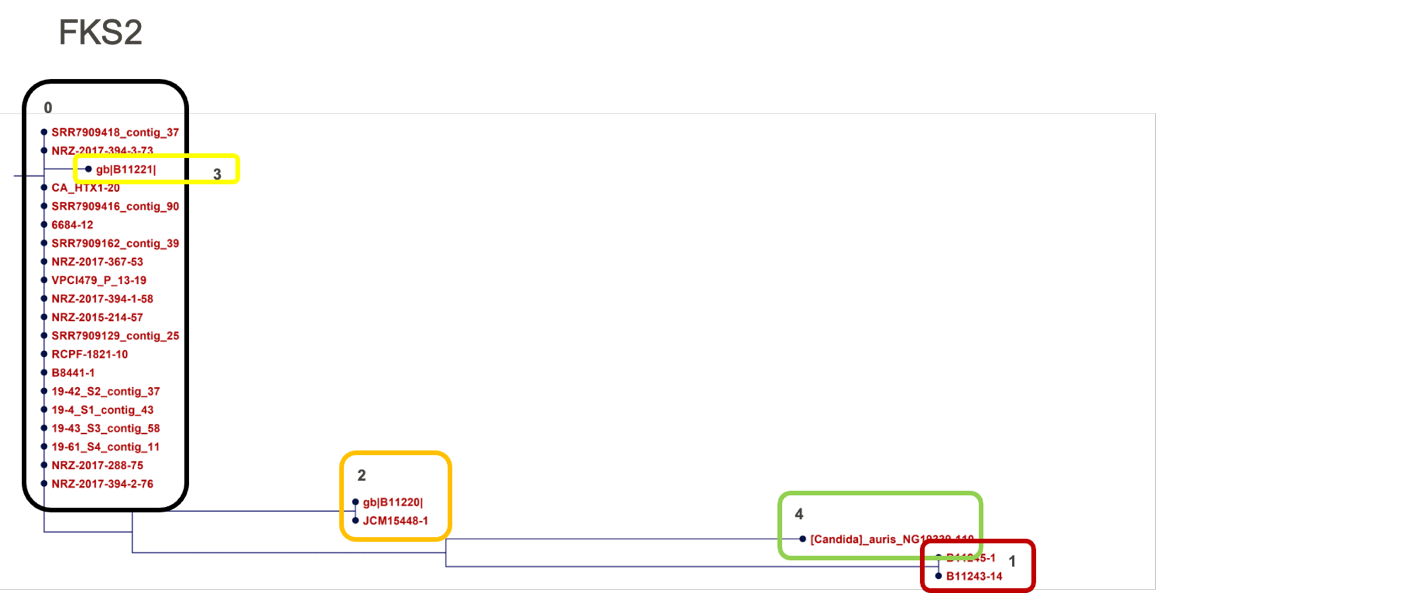

**Supplementary Figure 7-8**

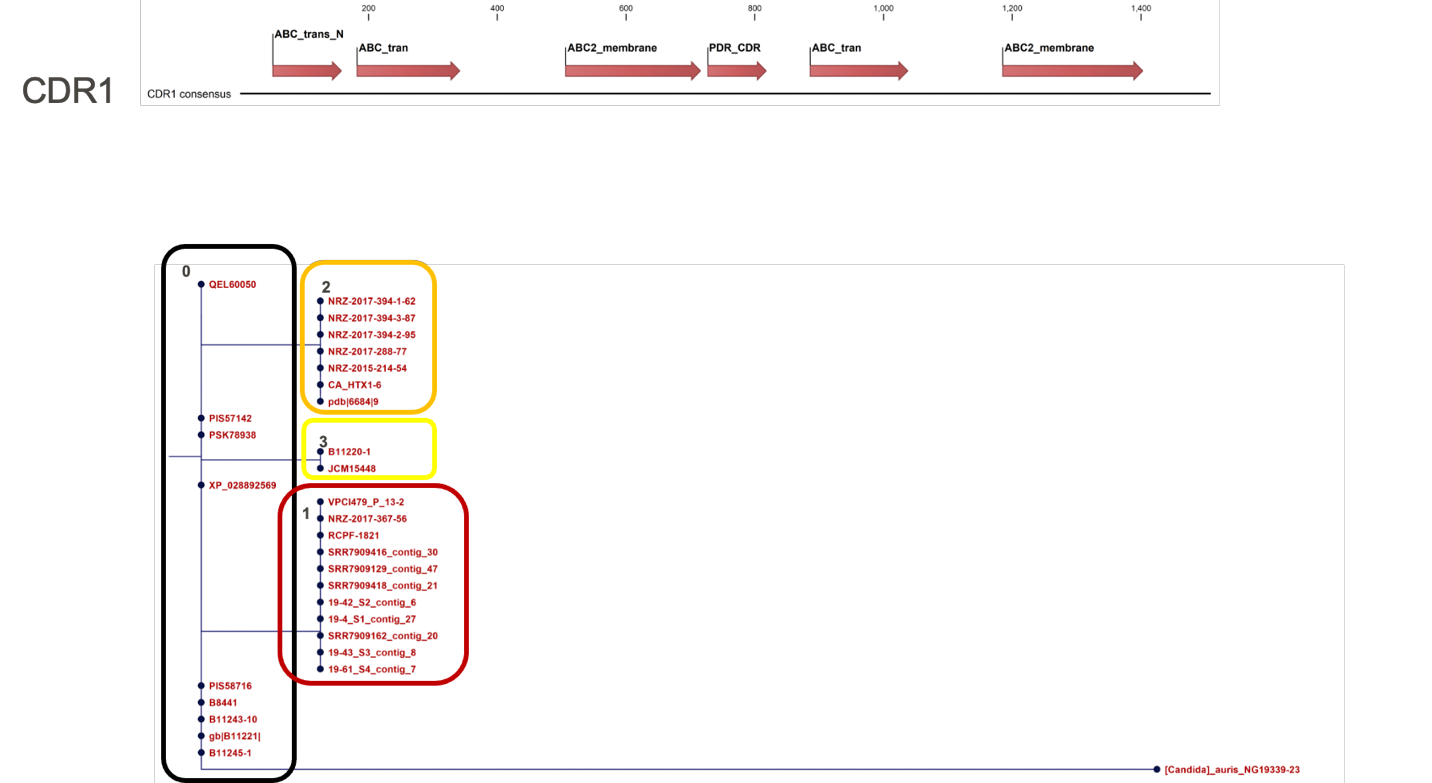

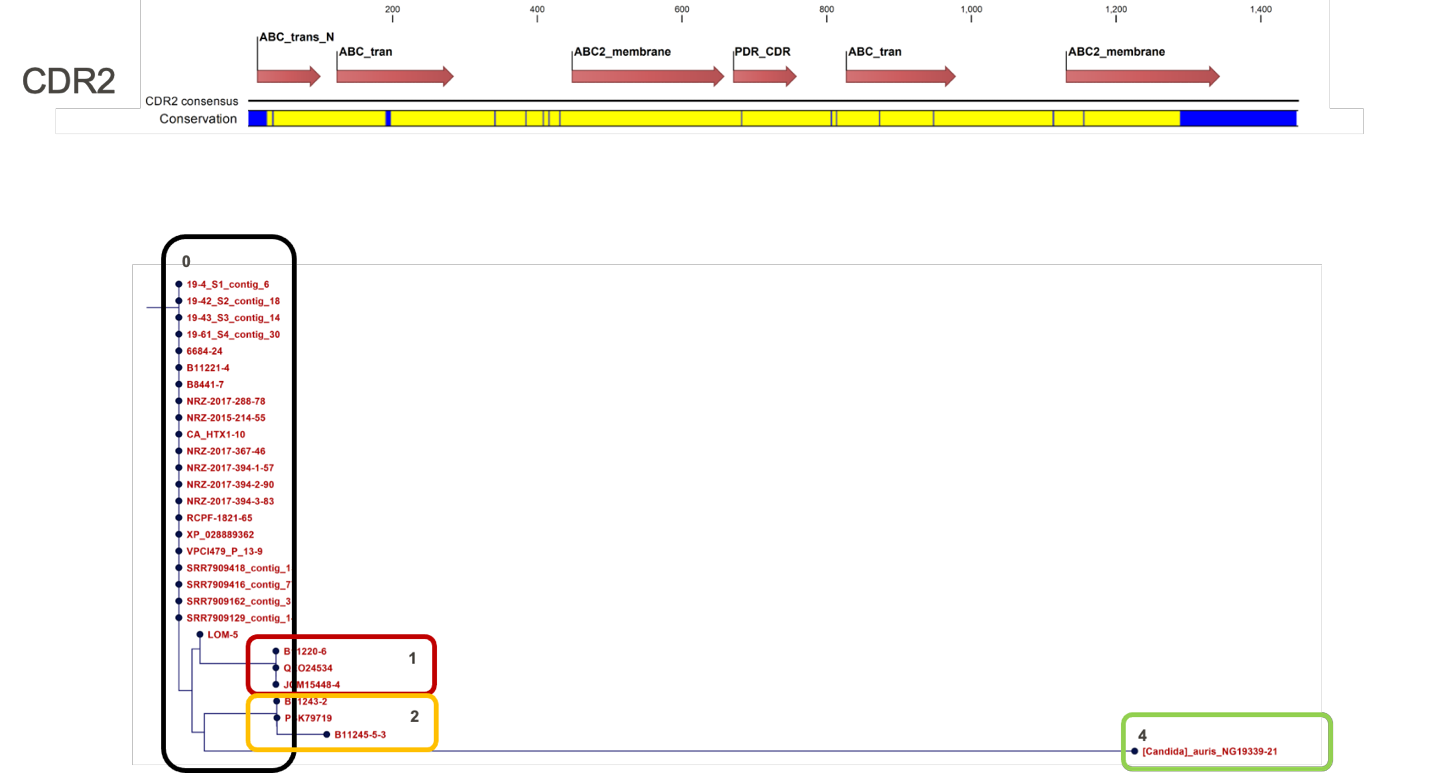

**Supplementary Figures 9-10**

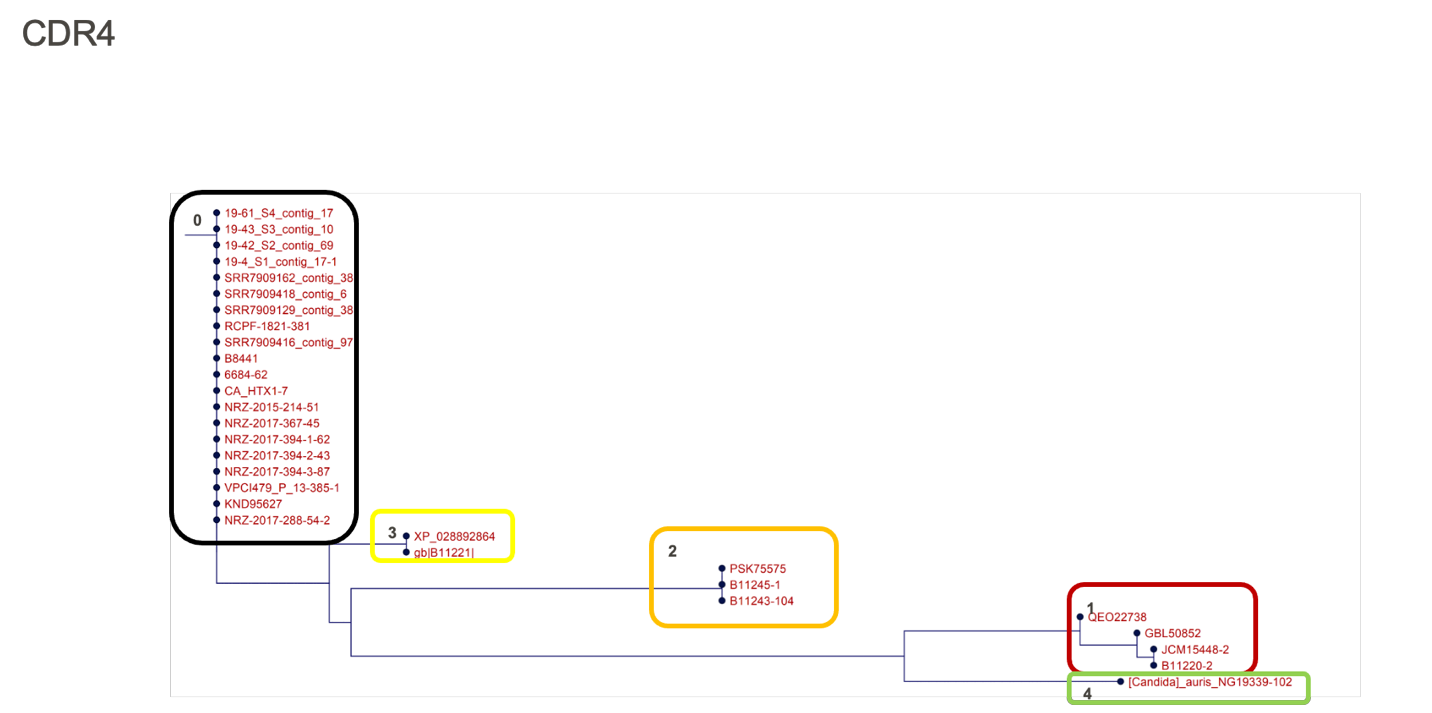

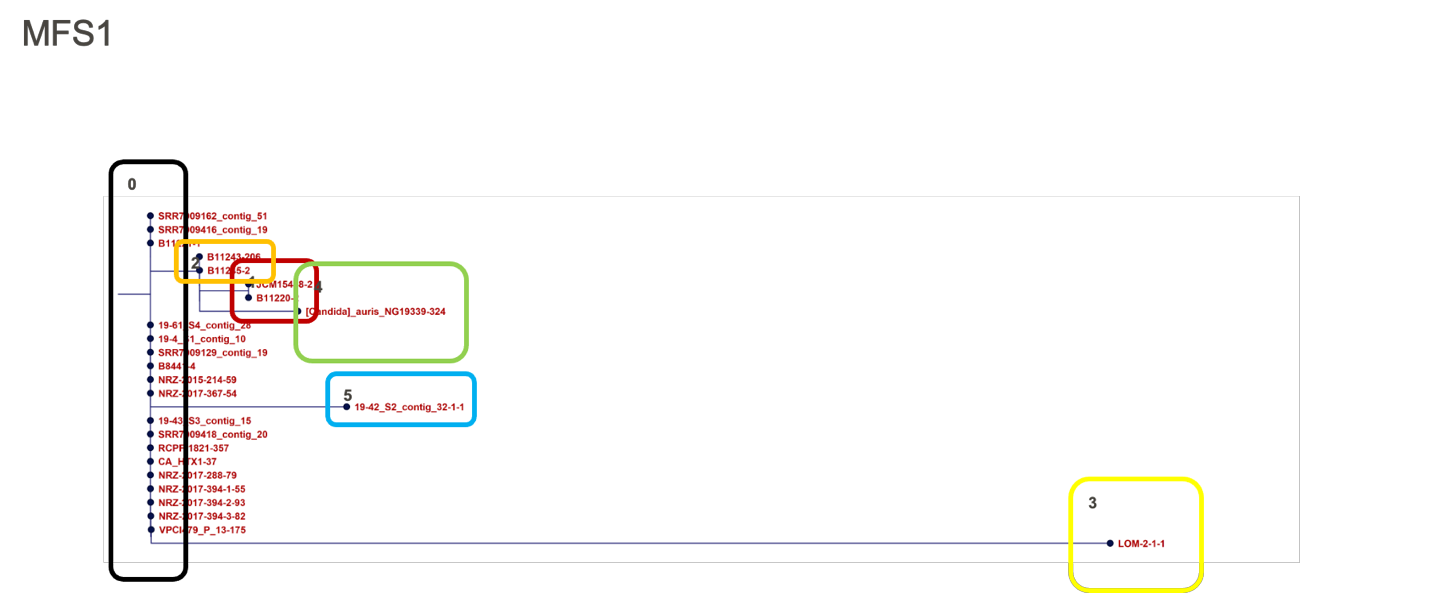

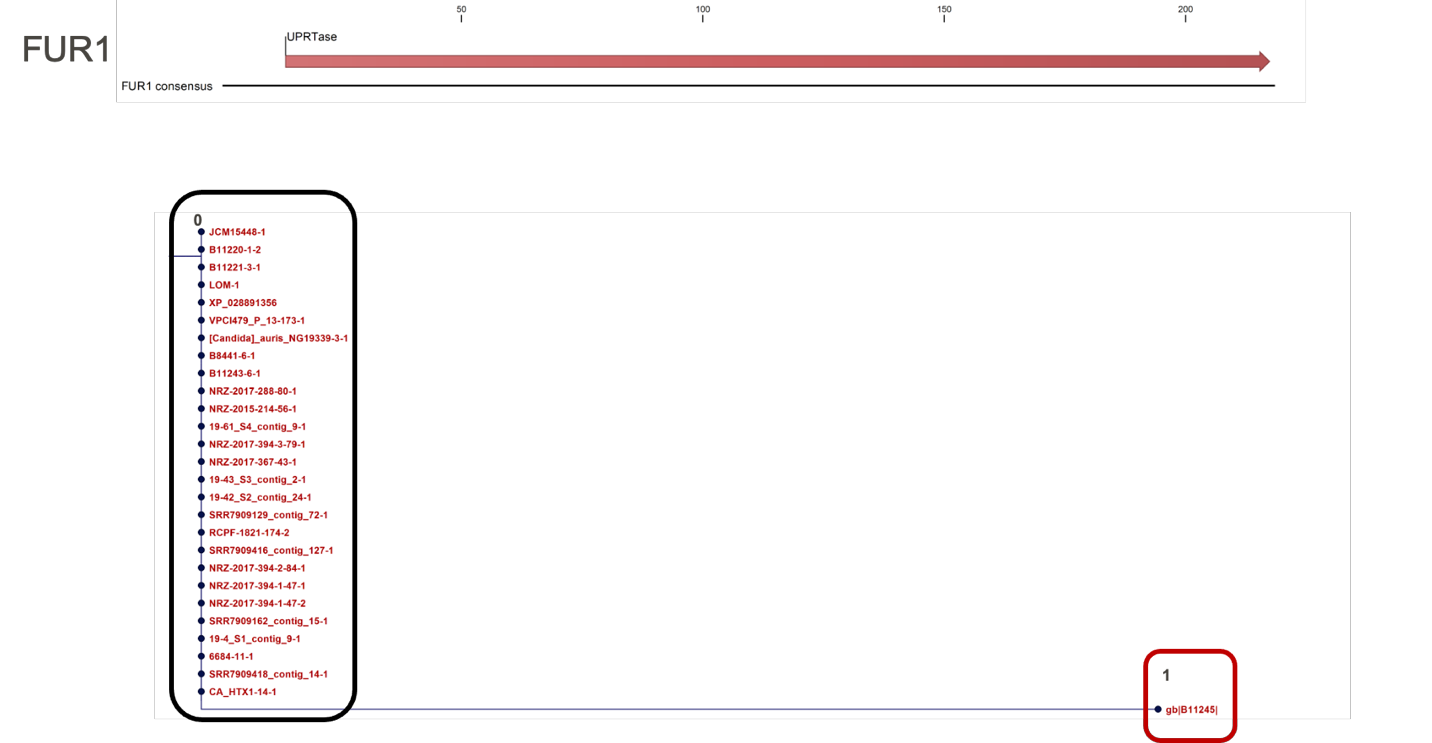
**Supplementary Figure 11-12**

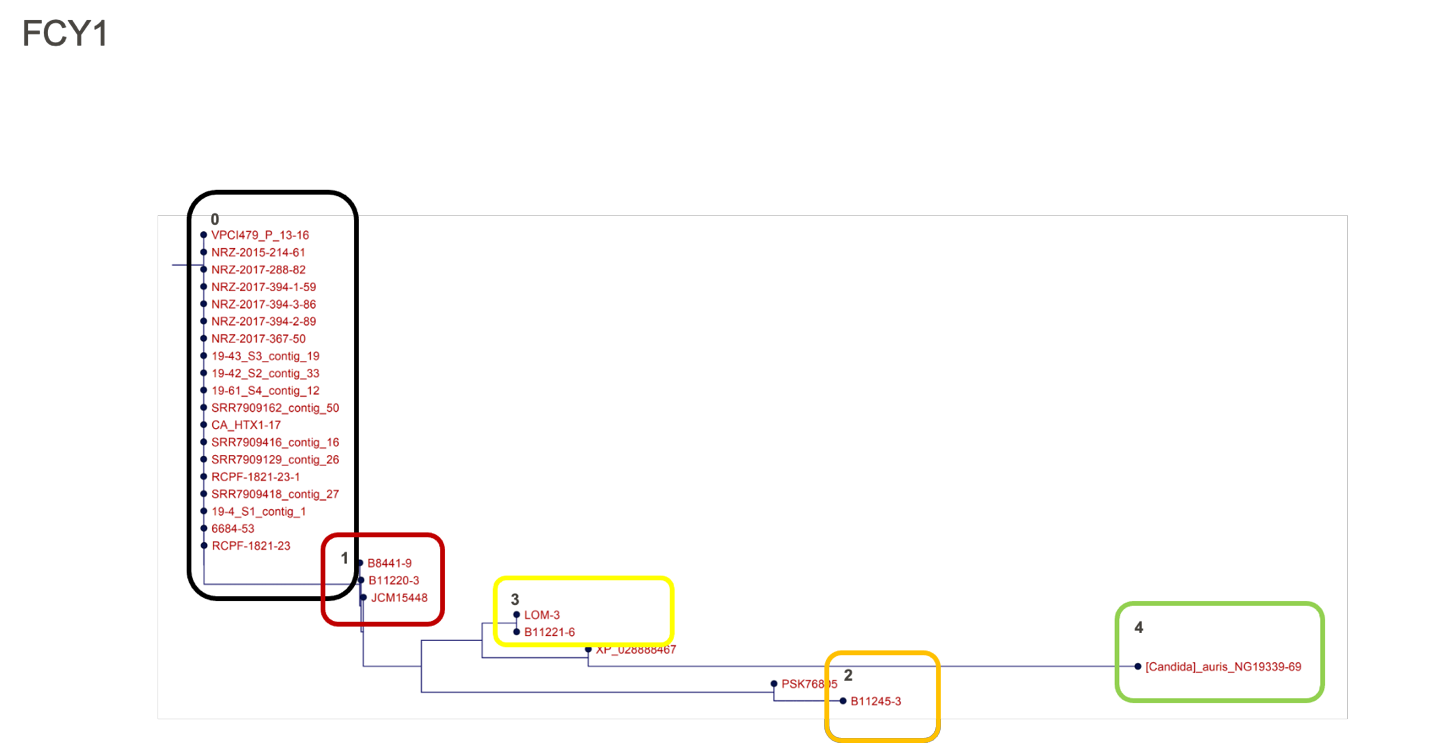

**Supplementary Figure 13-14**

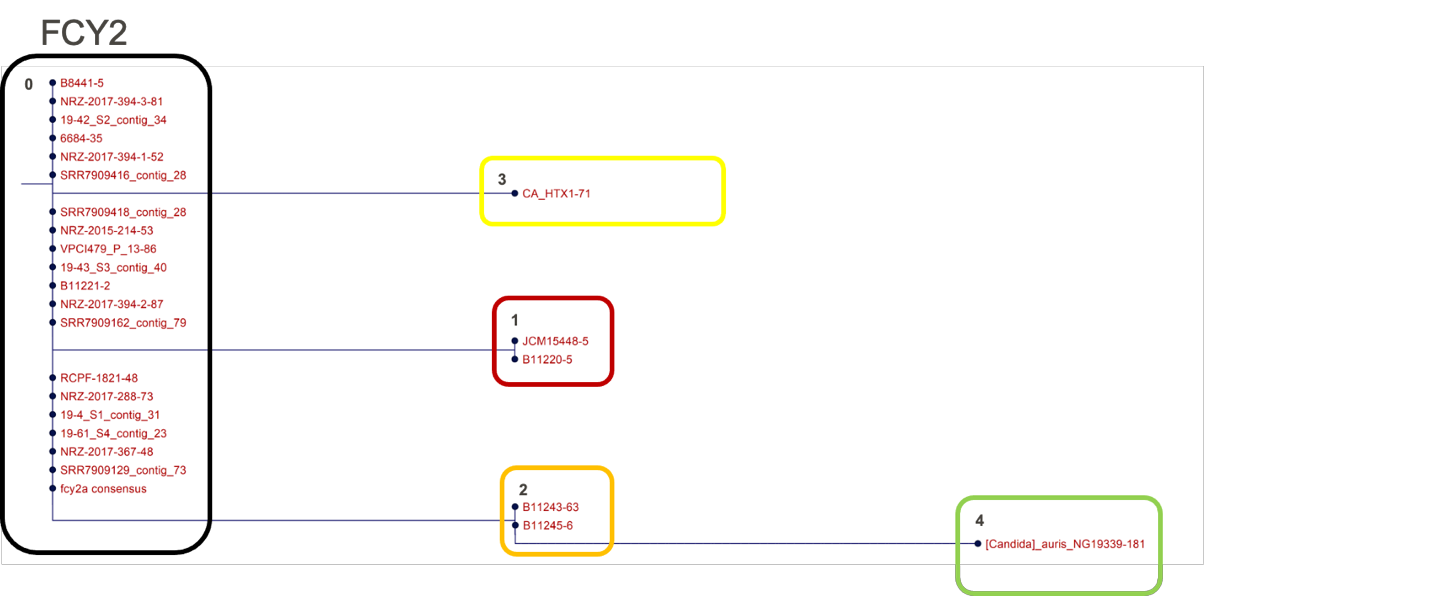

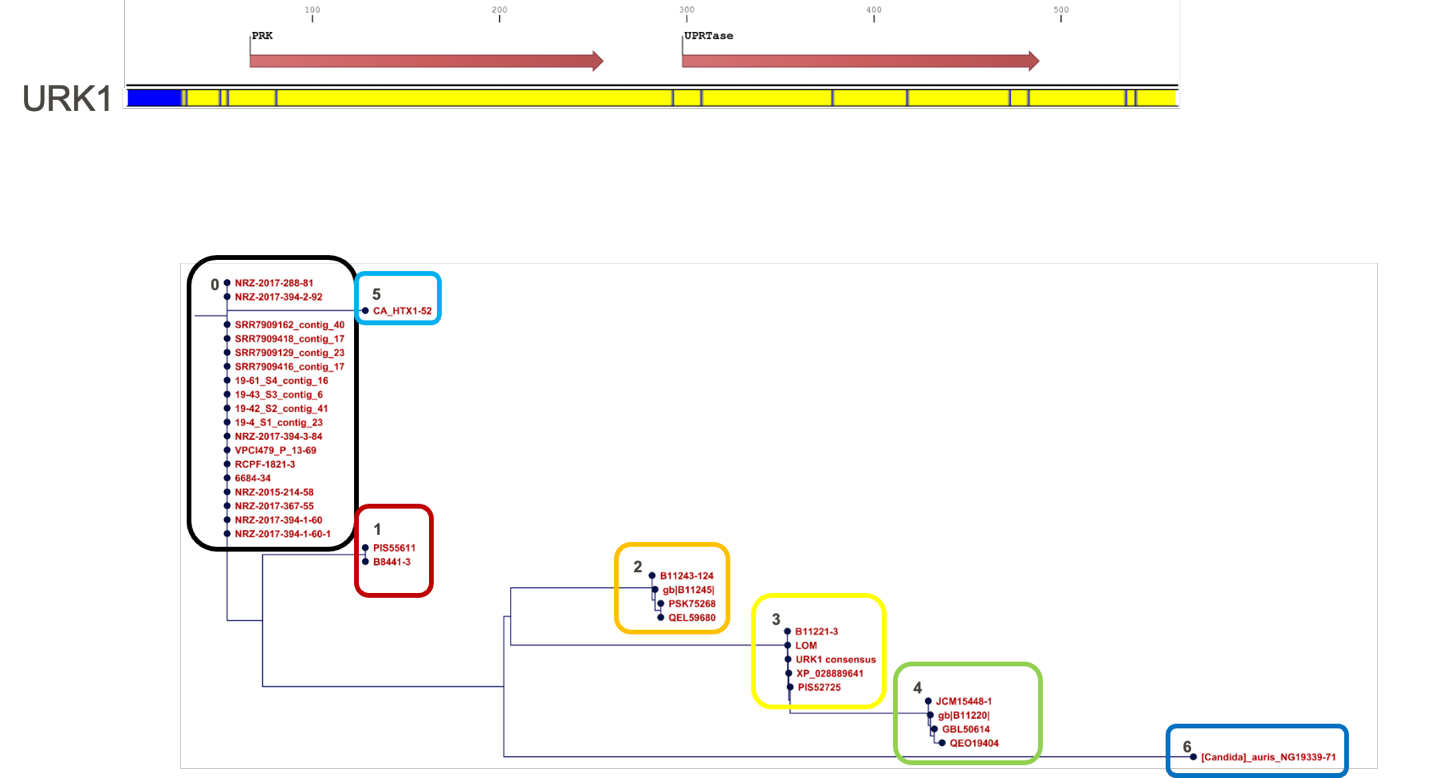
